# Perfusable 3D human urothelial model for real-time analysis of bacterial infection dynamics and therapeutic interventions

**DOI:** 10.1101/2025.11.12.687885

**Authors:** Amanzhol Kurmashev, Isabel Sorg, Julia Boos, Lara Grassi, Benjamin Sellner, Cinzia Fino, Mehmet Ugur Girgin, Antonia Müller, Steffi Klimke, Sarah Tschudin-Sutter, Andreas Hierlemann, Christoph Dehio

## Abstract

Urinary tract infections (UTIs) remain a major health burden, yet mechanistic studies are limited by the lack of experimental models that enable high spatiotemporal resolution tracking of infection dynamics, while recapitulating the stratified architecture of the bladder epithelium, urine tolerance and fluid dynamics. Here, we present a modular microphysiological platform integrating a fully stratified, urine-tolerant human urothelium cultured on standard transwell inserts within a custom-designed perfusion device compatible with live imaging. Urine flow enables real-time, high-resolution imaging of uropathogenic *Escherichia coli* (UPEC) infections under physiologically relevant conditions, including clearance of planktonic bacteria and nutrients replenishment, while retaining tissue-associated populations. This system revealed UPEC attachment via the type 1 fimbrial adhesin FimH and its inhibition by D-mannose treatment. Moreover, the platform captured L-form formation upon treatment with the frontline antibiotic fosfomycin and regrowth of walled bacteria following drug withdrawal. The platform further uncovered strain-specific lysis through bacteriophages in contrast to the activity of broad-spectrum antibiotics. In summary, this system constitutes a scalable platform with high predictive power for studying UTI pathogenesis and preclinical therapeutic testing.

## Introduction

Urinary tract infections (UTIs) are among the most common bacterial infections worldwide, with an estimated 400 million cases annually^1^. Uropathogenic *Escherichia coli* (UPEC), responsible for ⁓80% of cases, is therefore both a major cause for antibiotic prescription and a key driver of rising antimicrobial resistance (AMR)^2^. The clinical impact of resistant UPEC underscores the urgent need for improved strategies to study and treat these infections.

Progress has been hampered by limitations of current preclinical models. Animal models often fail to accurately reproduce human-relevant biology, which restricts translation to the clinics^3^. Similarly, standard in vitro assays assess compound efficacy using nutrient-rich growth media, such as Mueller-Hinton broth and agar, which do not mimic the complexity of a human urothelial tissue environment including urine exposure that are critical for host–pathogen interactions and infection dynamics^4^. As the primary interface between the urinary tract and its luminal environment, the urothelium plays a critical role in host defense through the exfoliation of infected cells and the production of a glycosaminoglycan (GAG) layer, which forms a protective biochemical interface that limits microbial adhesion^5^. Furthermore, urine composition, including its acidity and low nutrient content, represents a key factor of the local infection environment, influencing pathogen behavior and colonization dynamics^6, 7^.

To better recapitulate tissue complexity, several human-relevant bladder models have been developed, including ex vivo systems^8^, organoid cultures^9, 10^, and stratified transwell models^11^. Transwell-based systems have proven particularly powerful, enabling the formation of three-dimensional urine-tolerant human urothelial (3D-UHU) microtissues^11, 12^. However, conventional transwell systems lack dynamic aspects of bladder physiology, such as filling and voiding cycles, which affect bacterial adhesion and invasion^13^. In addition, their design hampers live imaging at single-cell resolution because of long working distances and refraction of light in the transwell membrane. Visualization of infection dynamics at high temporal and spatial resolution would, however, be invaluable to dissect host-pathogen interactions and uroepithelial responses in a physiologically relevant context.

Here, we present a microphysiological platform that integrates a stratified human urothelium with a urine flow system, compatible with high-resolution live-cell imaging. By culturing cells on the underside of standard transwell membranes, we generated tissues in an inverted orientation facing the lower reservoir that can be seamlessly integrated into a PDMS-free microfluidic device. This design allows for controlled perfusion of urine across the urothelial surface, recapitulating dynamic bladder conditions while maintaining tissue integrity. The use of high-resolution objectives and a glass-bottom interface enables live imaging of the infection process at the single-bacterial-cell level. After live experiments, the cultured tissue can be readily recovered for off-chip post-processing and histological or molecular analyses.

We validated the developed platform by monitoring the dynamics of UPEC infections and therapeutic interventions. The system revealed key roles of flagellar motility and type 1 fimbrial adhesin (FimH) in urothelial colonization, recapitulated antibiotic-induced formation of cell wall-deficient L-forms as observed in patient urine, and captured the outgrowth of rod-shaped bacteria upon drug withdrawal. It further enabled quantitative, time-resolved analysis of D-mannose-mediated competitive inhibition of adhesion and strain-specific bacteriophage activity. Together, these findings help to establish a physiologically relevant, imaging-compatible bladder model that overcomes a long-standing limitation in experimental UTI research. By combining human-like tissue architectures with a customizable flow device, the platform provides a powerful and scalable tool for preclinical studies of host-pathogen interactions and for the evaluation of precision antimicrobial therapies.

## Results

### Urine-tolerant human urothelial model for live-cell imaging

To enable real-time imaging and improve experimental throughput, we established an inverted 3D human urothelial transwell model adapted to a 24-well plate format (Fig. 1a). HBLAK human bladder progenitor cells were seeded on the underside of transwell insert polycarbonate membranes after the inserts were flipped (day 0). Following overnight incubation, the inserts were flipped back to the upright position (day 1) and expanded in 2D growth medium (CnT-Prime) at a liquid-liquid interface (LLI) until confluence (day 4).

**Figure 1.**
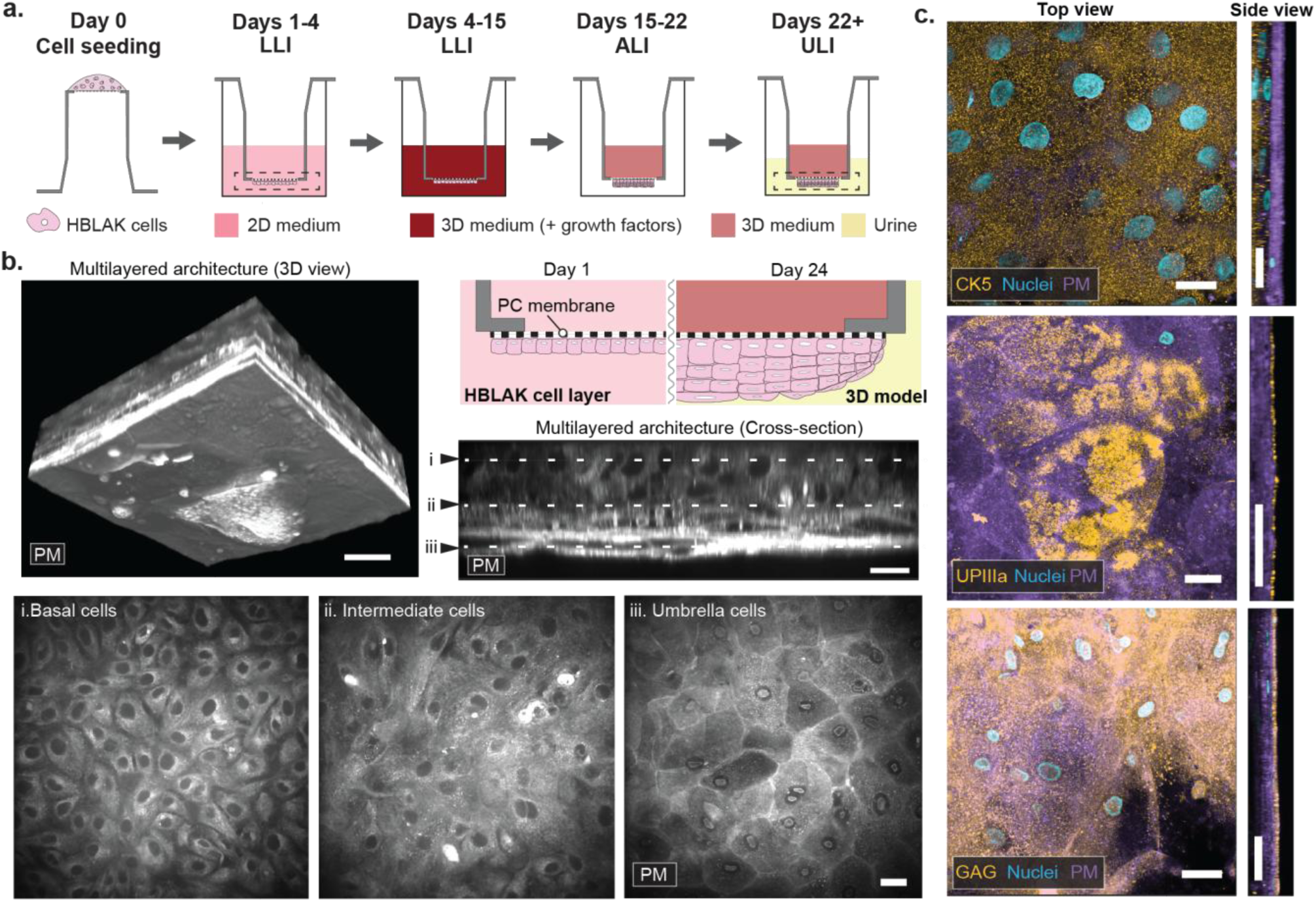
Generation of functionally stratified bladder tissue for live cell imaging. **a,** Schematic representation of sequential steps in tissue formation using the inverted culture approach. Human bladder epithelial cells (HBLAK) were seeded on the underside of transwell inserts and maintained under three culture conditions: liquid–liquid interface (LLI), air-liquid interface (ALI), or urine–liquid interface (ULI). **b,** 3D reconstruction and cross-sectional view of live stratified bladder tissue cultured on an inverted polycarbonate (PC) transwell insert. Plasma membranes were labeled with CellMask (grey). Basal cells (i), intermediate cells (iii), and umbrella cells (ii) are shown. Scale bars: 20 µm. **c,** Immunofluorescence analysis of cell-type-specific and barrier-integrity markers in 3D bladder tissue: cytokeratin-5 (CK-5, basal cells, yellow), uroplakin IIIa (UPIIIa, umbrella cells, yellow), and heparan sulfate proteoglycan 2 (glycosaminoglycan (GAG) layer, yellow). The nuclei were stained with DAPI (cyan) and the plasma membranes (PM) with CellMask (purple). Side views are shown below. Scale bars, 20 µm.

The cultures were then switched to 3D differentiation medium (3D-Prime) supplemented with growth factors to induce stratification and basal cell proliferation. In contrast to the upright 3D-UHU protocol^11^, where urine is introduced on day 3, our model was maintained in supplemented differentiation medium until day 15. At this point, an air-liquid interface (ALI) was established by adding 3D differentiation medium to the upper chamber only. The ALI phase proved essential for tissue integrity, as premature urine exposure or omission of the ALI phase resulted in epithelial breakdown (data not shown). CellMask™ Plasma Membrane Deep Red (CellMask) staining enabled live tracking of tissue morphology throughout differentiation.

From day 22 onward, cultures were transitioned to a urine-liquid interface (ULI), with pooled human urine in the lower chamber and 3D differentiation medium in the upper chamber. Confocal microscopy on day 27 revealed the formation of a stratified epithelium composed of 6-8 layers, including basal, intermediate, and umbrella cells (Fig. 1b). The basal cells were located adjacent to the transwell membrane, while the umbrella cells formed the apical surface directly exposed to urine. Immunocytochemistry confirmed urothelial differentiation on day 25: cytokeratin 5 marked basal cells, uroplakin IIIa labeled umbrella cells, and perlecan (heparan sulfate proteoglycan 2) stained the glycosaminoglycan (GAG)-rich surface layer (Fig. 1c). The expression of the tight junction protein ZO-1, together with the increase in TEER values between days 15 and 22 (Extended Data Fig. 1a,b), indicated barrier maturation, coinciding with the emergence of umbrella cells observed by live cell imaging.

### Microfluidics-based recapitulation of urine flow

A defining feature of urinary tract physiology is the unidirectional flow of urine across the apical urothelium. To reproduce this microenvironment under physiologically relevant infection conditions, we integrated our 3D urothelial model into a poly(methyl methacrylate) (PMMA)-based microfluidic platform that accommodates four transwell inserts (Fig. 2a). Each insert was mounted into a microfluidic unit and secured with an O-ring that expanded upon insertion to form a leak-proof seal for urine perfusion (Fig. 2a,b). The platform design is readily adaptable to different commercial inserts by adjusting well dimensions and O-ring sizes (Supplementary Fig. 1a,b). Microfabricated pillars at the base of each well positioned the insert 300 µm above the glass coverslip (Fig. 2b), enabling high-resolution live imaging (up to 40× magnification) during perfusion. The microchannel beneath the insert incorporated phase guides (Fig. 2a) to facilitate bubble-free filling and to ensure uniform urine flow and shear stress across the tissue (Supplementary Fig. 1c,d).

**Figure 2.**
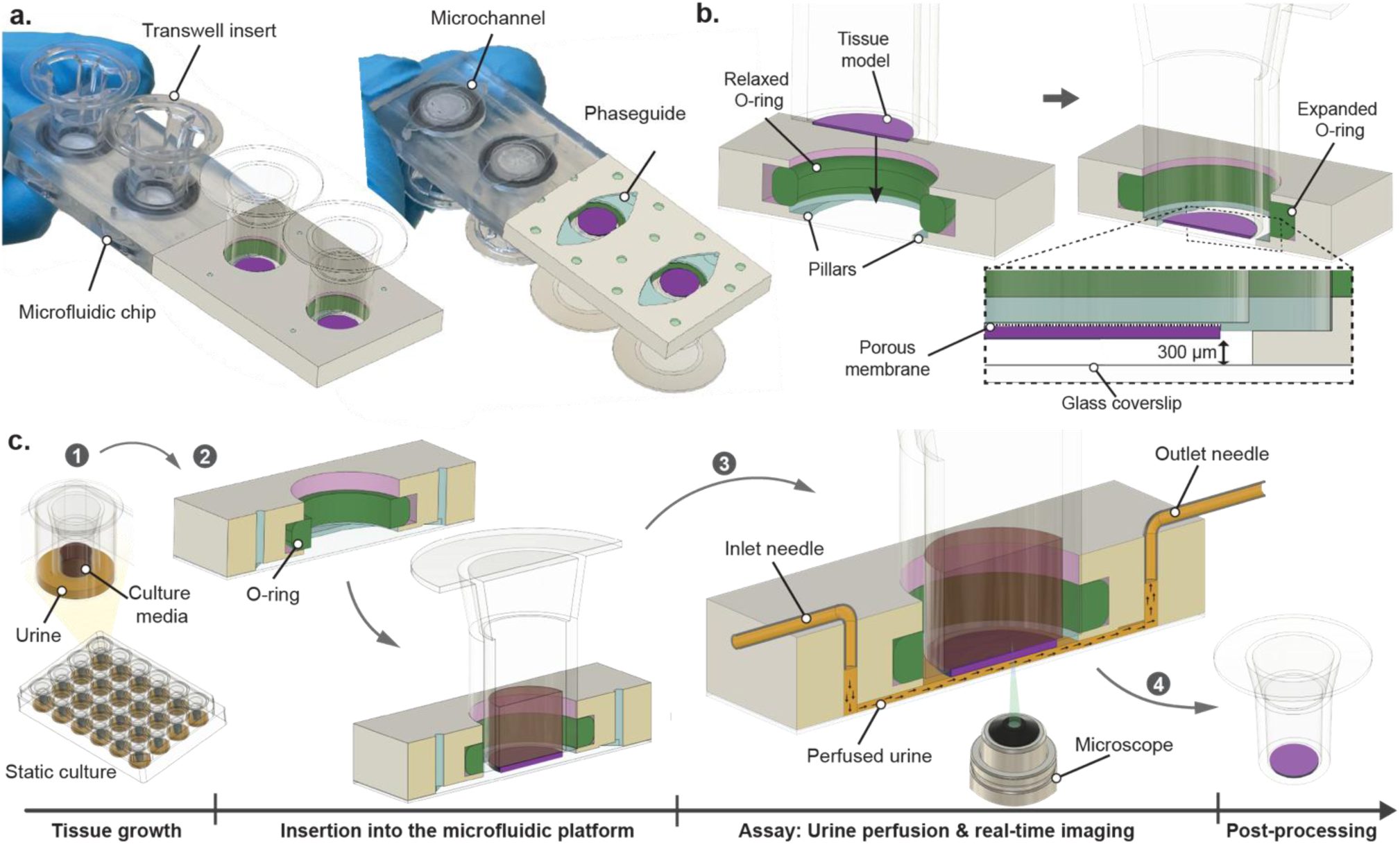
Microfluidic platform and experimental setup. **a,** Schematic of the custom-designed microfluidic chip. **b,** Transwell inserts were mounted with a relaxed O-ring that expanded upon insertion to form a leak-proof seal for urine perfusion. Two lateral plastic pillars supported the insert and defined a 300-µm imaging distance between the tissue and the glass bottom. **c,** Overview of the experimental workflow to assess tissue stability during transwell insertion (steps 1–2) and urine perfusion (step 3), followed by live imaging and post hoc analysis (step 4).

The modular design allowed tissues to be grown under static conditions, permitting quality control and selection of optimal inserts prior to integration into the perfusion system for infection assays with high-resolution imaging. Following experiments, inserts could be easily recovered for endpoint analyses, including TEER measurements and immunofluorescence stainings (Figure 2c).

To assess the effect of urine flow on tissue stability, urothelial cultures were maintained under static conditions or perfused at 100 µl min⁻¹ for 6 h. Live cell staining revealed intact umbrella cell layers under both conditions (Supplementary Fig. 2a). TEER measurements revealed a modest decline for both conditions, with a greater reduction under flow (Supplementary Fig. 2b), likely reflecting minor edge damage incurred during transwell retrieval.

### Enhanced model relevance via orientation and urinary flow

Beyond adherence, bacterial motility is a key virulence determinant in urinary tract infections^14^. To examine how tissue orientation and urine flow influence infection dynamics of motile wild type (WT) and non-motile Δ*fliC* UPEC, we compared the conventional upright transwell model (mimicked by flipping the inverted tissue), where under static conditions bacteria settle onto the tissue by gravity, with our inverted setup operated under static and perfused conditions (Fig. 3a). Adhesion of the Δ*fliC* mutant was significantly reduced in both, static and perfused, inverted models compared to the motile wild type (Fig. 3b,c), underscoring the importance of active motility for successful surface colonization when passive sedimentation is prevented and thereby reflecting a more physiological scenario during ascending infection in the urinary tract.

**Figure 3.**
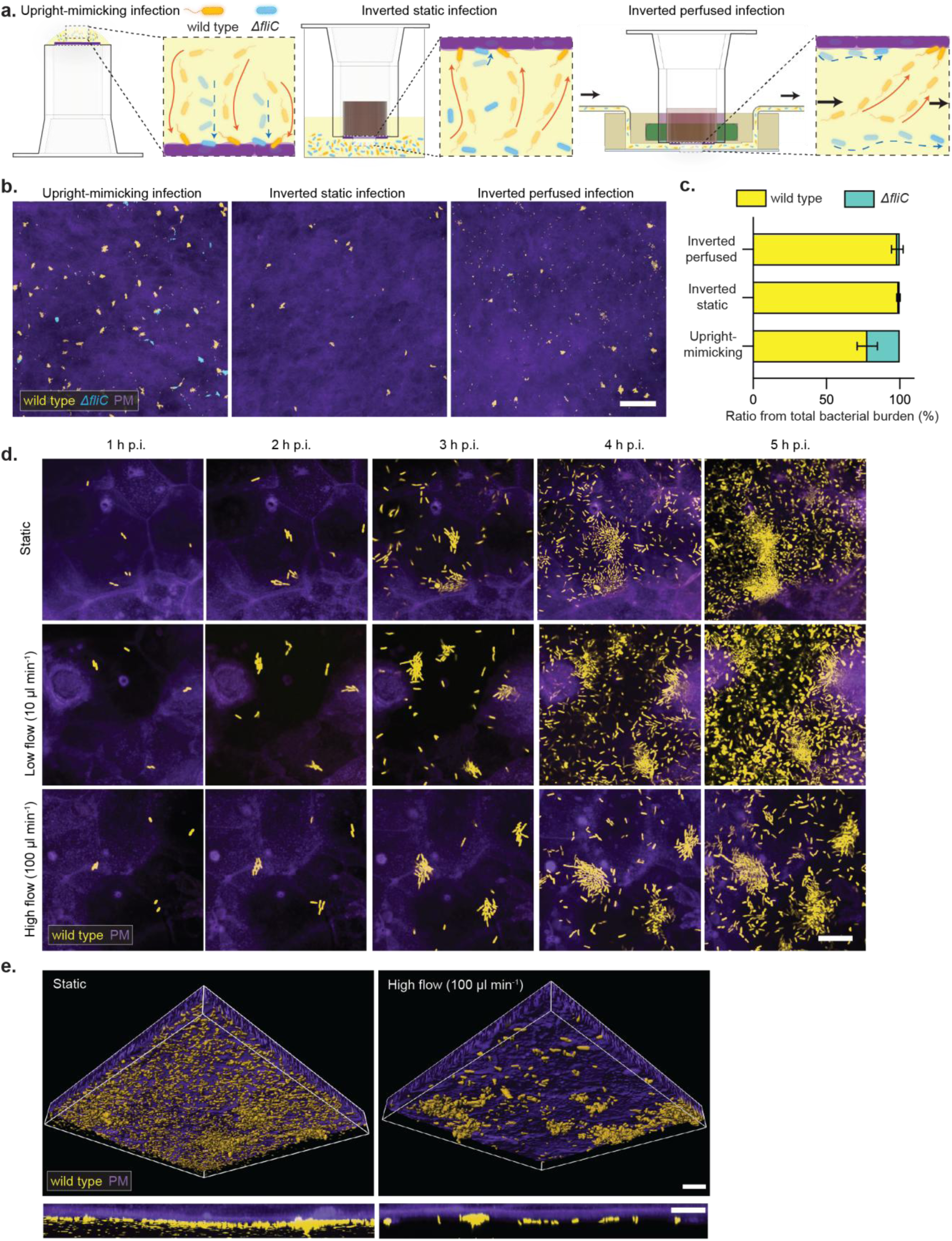
Effect of urinary flow and tissue orientation on UPEC infection. **a,** Schematic of infection experiments using the motile UPEC strain CFT073 expressing mScarlet3 (wild type, yellow) and the isogenic non-motile mutant Δ*fliC* expressing mGreenLantern (cyan) in upright-mimicking, inverted static, and inverted perfused transwells. **b,** Representative fluorescence images of tissues infected for 3 hours with a 1:1 mixture of wild type (yellow) and Δ*fliC* (cyan) CFT073 under the indicated conditions. Scale bar, 100 µm. **c,** Quantification of surface-associated bacteria 3 h post infection (h p.i.). For each condition, 25 images per insert were acquired at 25× magnification from at least two inserts per condition. The WT/Δ*fliC* ratio is shown for the respective infection setups. Data represent mean ± s.d. from *n* = 3 biologically independent experiments. **d,** On-chip infection with wild type CFT073 under different flow regimes. The bacteria were perfused for 30 min at 100 µl min⁻¹, followed by a 30-min wash with sterile urine at the same flow rate. At 1 h p.i., flow was stopped (static) or maintained at 10 µl min⁻¹ or 100 µl min⁻¹ until 5 h p.i. Host cell plasma membranes (PM) were stained with CellMask (purple). Representative live-cell fluorescence images are shown for each condition (*n* = 3 biologically independent experiments). Scale bars, 20 µm. **e,** 3D reconstructions of the host-pathogen interface at 4 h p.i. wild type bacteria (yellow) localize on the tissue surface, shown in a tilted 3D view (top) and the corresponding side view (bottom). Representative images for the conditions indicated in (d). Scale bar: 20 µm.

Next, we assessed the impact of urine flow, a critical in vivo factor that removes planktonic bacteria and replenishes nutrients, which maintain uropathogens in an active growth state. Real-time monitoring under static conditions, low (10 µl min⁻¹) and high (100 µl min⁻¹) flow conditions (Fig. 3d) revealed distinct stages of UPEC infection: (i) initial attachment (0–1 h post infection, hpi), (ii) microcolony formation (1–4 hpi), and (iii) widespread surface colonization (4–5 hpi). At 4 hpi, static and low-flow conditions exhibited extensive planktonic overgrowth of the urothelium, whereas high-flow (100 µl min⁻¹) effectively removed planktonic and loosely attached bacteria, but preserved firmly attached bacteria (Fig. 3d,e), likely reflecting the bacterial clearance occurring in vivo during bladder filling and voiding cycles. Together, these findings emphasize that the incorporation of physiological urine flow and tissue orientation markedly enhances the biological relevance of in vitro UTI models.

### Model benchmarking against clinical and ex vivo observations of fosfomycin-induced L-form formation and regrowth of rod-shaped bacteria upon drug removal

UPEC can transiently adopt cell wall-deficient (L-form) states under antibiotic stress, allowing bacteria to survive antibacterial treatment for prolonged time. A study identified L-forms in fresh urine samples from elderly patients with recurrent UTIs (rUTIs) receiving the cell wall-targeting antibiotic fosfomycin.^15^ In ex vivo urine samples, UPEC readily transitioned from rod-shaped to the L-form morphologies during fosfomycin treatment and seemingly reverted to rods after drug removal, with L-form formation and survival supported by the osmoprotective properties of urine.^15^

To benchmark our urothelial tissue model against these clinical and ex vivo findings, we reproduced the reported conditions under controlled urine flow (Fig. 4a). Urothelial tissue infected with the UPEC strain CFT073 was exposed to a continuous flow of urine containing 400 mg l⁻¹ of fosfomycin, a concentration representative of urinary levels during several hours after administration of a single dose.^16^ Within five hours of exposure, most rod-shaped bacteria were lysed or converted to L-forms, with higher urine osmolarity providing greater protection against osmotic lysis (Fig. 4b). To capture the rare events of rod-shaped regrowth upon drug withdrawal, we imaged z-stacks of multiple regions of the microtissue at 30 min intervals. Using this approach, we recorded two instances of rod-shaped regrowth in a single high-osmolarity urine sample (Fig. 4c). However, this temporal resolution was insufficient to identify the initial bacterial cell that resumed growth via lineage tracing. To characterize such rare events in greater detail, we reduced system complexity by using axenic conditions in CellASIC microfluidic chambers operated under controlled diffusive flow, enabling the application of identical exposure profiles across different urine samples.^17^ High-resolution imaging at 2 min intervals confirmed the same osmolarity-dependent pattern of L-form conversion and survival, yet regrowth of rod-shaped cells was observed in all urines across the spectrum of osmolarities (Supplementary Table S1). A detailed analysis of the recordings identified only a single L-to-rod reversion event (Supplementary Movie S1), whereas the regrowth in other cases originated from non-dividing rods with diminished fluorescence expression, indicating metabolic quiescence that spontaneously resumed growth and fluorescence expression after drug removal (Supplementary Movie S2). The presence and growth of these initially non-dividing rods were largely unaffected by urine osmolarity, indicating no apparent correlation between osmolarity and regrowth.

**Figure 4.**
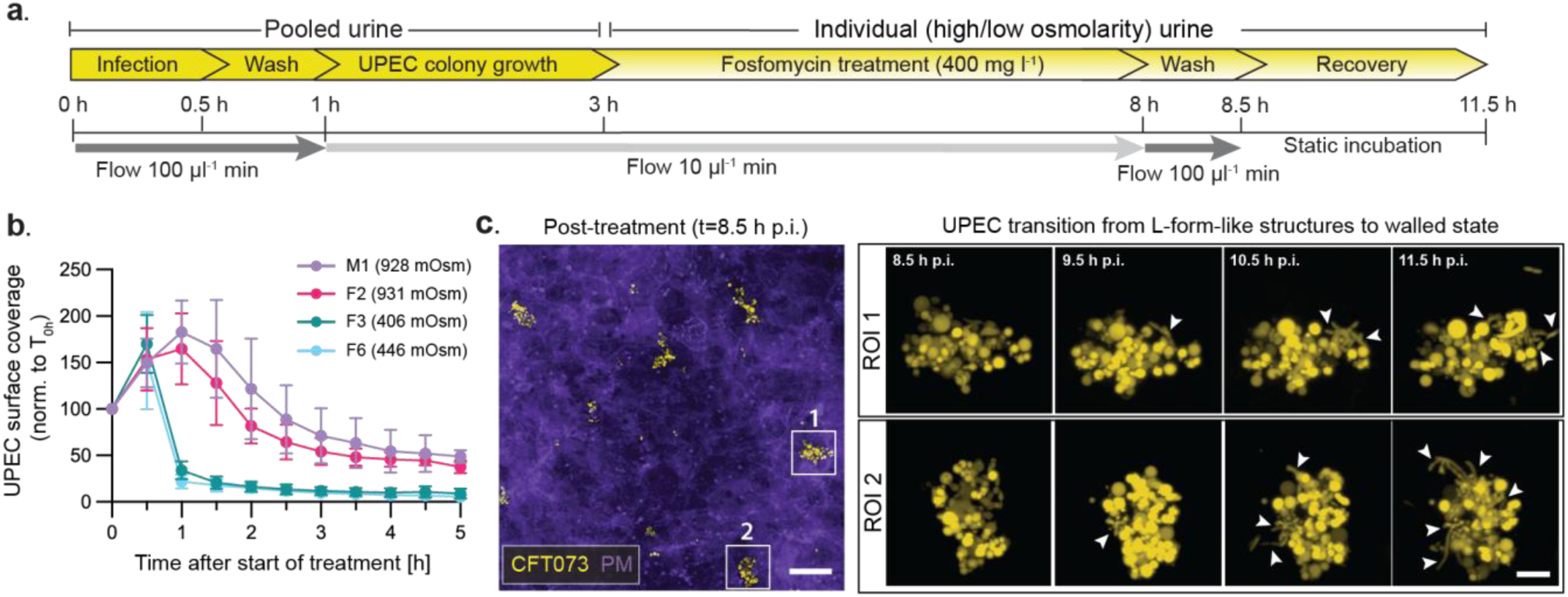
Effect of urine osmolarity on L-form stability induced by fosfomycin, and real-time visualization of UPEC regrowth upon drug withdrawal. **a,** Experimental workflow and timeline illustrating UPEC infection by strain CFT073, fosfomycin exposure, and subsequent recovery following drug removal. **b,** Temporal dynamics of UPEC surface coverage on the urothelial tissue during fosfomycin treatment in urine samples of high (M1, F2) or low (F3, F6) osmolarity. UPEC surface coverage was measured and normalized to baseline (T0 = 100%). Data are shown as mean ± s.d. from three independent experiments (n = 3). **c,** Left: Representative fluorescence image of infected urothelial tissue after fosfomycin exposure, showing residual UPEC L-form microcolonies. Scale bar, 50 µm. Right: Zoomed-in regions of interest (ROIs) showing time-lapse sequence of some L-form cells reverting to rod-shaped morphology during regrowth after removal of fosfomycin. Scale bar, 5 µm.

Finally, batch-culture time-kill experiments in 96-well microtiter plates demonstrated that elevated urine osmolarity significantly enhanced bacterial survival during fosfomycin exposure, corroborating survival rates obtained with the tissue model (Fig. 4b, Supplementary Fig. S3).

Together, these results show that our urothelial tissue model faithfully reproduces clinically observed L-form formation under fosfomycin treatment and suggests that post-antibiotic regrowth predominantly arises from metabolically dormant, non-dividing rods, while also L-form reversion may occur at low frequency. This mechanistic insight provides a new framework for understanding recurrent UTI relapses and suggests that targeting non-dividing populations may be critical for improved therapeutic outcomes.

### D-mannose attenuates FimH-mediated UPEC adhesion under flow

FimH, the adhesin located at the tip of type 1 pili, is a key UPEC virulence factor that binds mannosylated uroplakins to mediate host attachment and promote microcolony formation, a process competitively inhibited by D-mannose (Fig. 5a).^18^ To validate FimH-mediated adhesion in our microphysiological platform, human bladder tissues were co-infected with wild type UPEC expressing mScarlet3 (wild type, yellow) and an isogenic FimH-deficient mutant expressing mGreenLantern (Δ*fimH,* cyan) for 3 h to allow initial microcolony establishment, followed by perfusion with urine containing 0%, 1%, or 7% D-mannose (Fig. 5b). Live-cell imaging revealed that, in untreated controls, wild type formed dense microcolonies on the urothelial surface within 5 h post-infection, whereas the Δ*fimH* mutant exhibited only a sparse, unstructured attachment without interbacterial organization (Fig. 5c). Perfusion with D-mannose markedly disrupted wild type adhesion and microcolony formation, confirming that FimH-mannose interactions were involved in early UPEC colonization (Fig. 5c). Time-lapse imaging also showed flow-mediated detachment of preformed wild type clusters upon mannose exposure, illustrating the real-time effect of competitive inhibition of FimH-mediated binding (Fig. 5c, Supplementary Movie 3). Quantification confirmed a strong decrease of wild type levels under mannose-treatment conditions (Fig. 5d; Supplementary Fig. S4a), while Δ*fimH* mutant levels were consistently low across all treatments.

**Figure 5.**
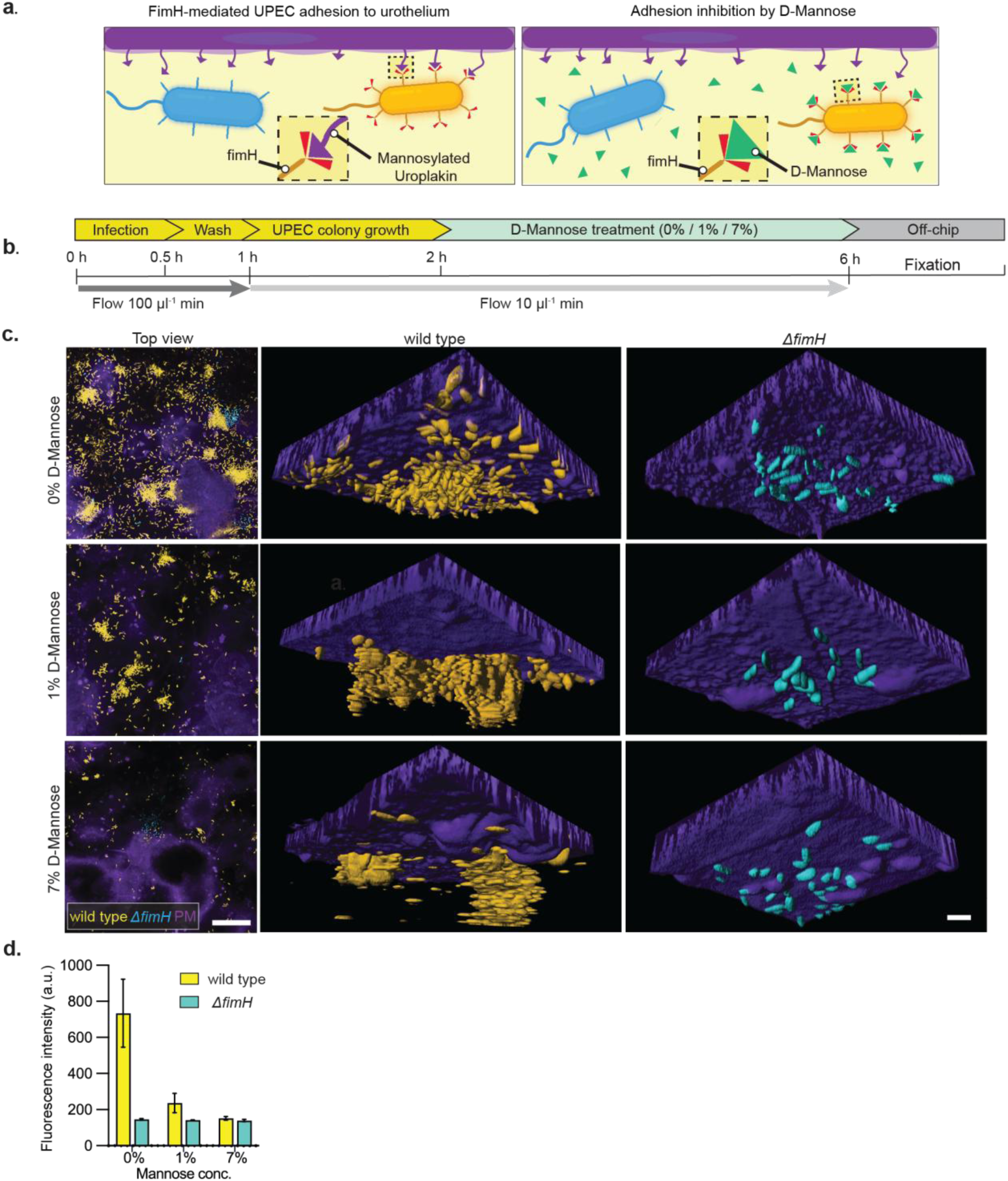
D-mannose attenuates FimH-mediated UPEC colonization of the urothelium under urine flow. **a**, Schematic illustration of FimH-dependent binding of UPEC strain CFT073 expressing mScarlet3 (WT, yellow) to mannosylated uroplakin receptors on umbrella cells of the urothelium (left). Competitive inhibition by D-mannose prevents FimH binding (right), whereas the isogenic FimH-deficient mutant Δ*fimH* expressing mGreenLantern (cyan) fails to adhere. **b**, Experimental workflow and corresponding timeline. **c,** Representative live-cell imaging and 3D reconstructions of urothelial tissue infected with a 1:1 mixture of wild type (orange) and Δ*fimH* (cyan) under continuous urine flow, with 0%, 1%, or 7% D-mannose. D-mannose was administered from 2 to 5 h after infection (hpi). Left panels: live-cell images at 5 hpi (40× objective). Scale bar, 50 µm; Middle and right: 3D reconstructions showing surface-associated wild type and Δ*fimH* bacteria. Urothelial plasma membranes (PM) were visualized using CellMask (purple). Scale bar, 5 µm. **d**, Quantification of bacterial surface colonization, based on mean fluorescence intensity across the urothelial surface, in the absence or presence of the indicated concentrations of D-mannose (applied from 2 to 5 h post-infection, hpi). Data represent mean ± s.d. from *n* = 3 biologically independent experiments.

To analyze the contribution of the tissue microenvironment to FimH-independent attachment, we perfused urine supplemented with inert carboxylated nanoparticles. These particles adhered at low frequency to the epithelial surface (Supplementary Fig. 4b), suggesting that, in the absence of FimH-dependent interactions, a fraction of bacterial binding still occurs through FimH-independent mechanisms.

### The urothelial model reveals the elimination of UPEC strains by strain-specific phages

As the threat of AMR is increasing, alternatives to antibiotics such as phage therapy gain more and more interest. Unlike antibiotics phages are highly strain specific with little impact on other microorganisms and a reduced risk of broad spectrum-resistance.^19^ We tested two strain-specific phages targeting clinically relevant UPEC strains (Supplementary Figs. 5a, b). The bacterial hosts were differentially labeled with fluorescent proteins (CFT073 shown in yellow, UTI89 shown in cyan) and co-infected in our urothelial tissue model (Fig. 6a). After both strains had established widespread surface colonization (4.5 h post-infection), the tissues were exposed for 90 min to urine flow with the corresponding strain-specific phages or, as a control, to the combination of amoxicillin and clavulanic acid, a first-line broad-spectrum antibiotic (Fig. 6b).

**Figure 6.**
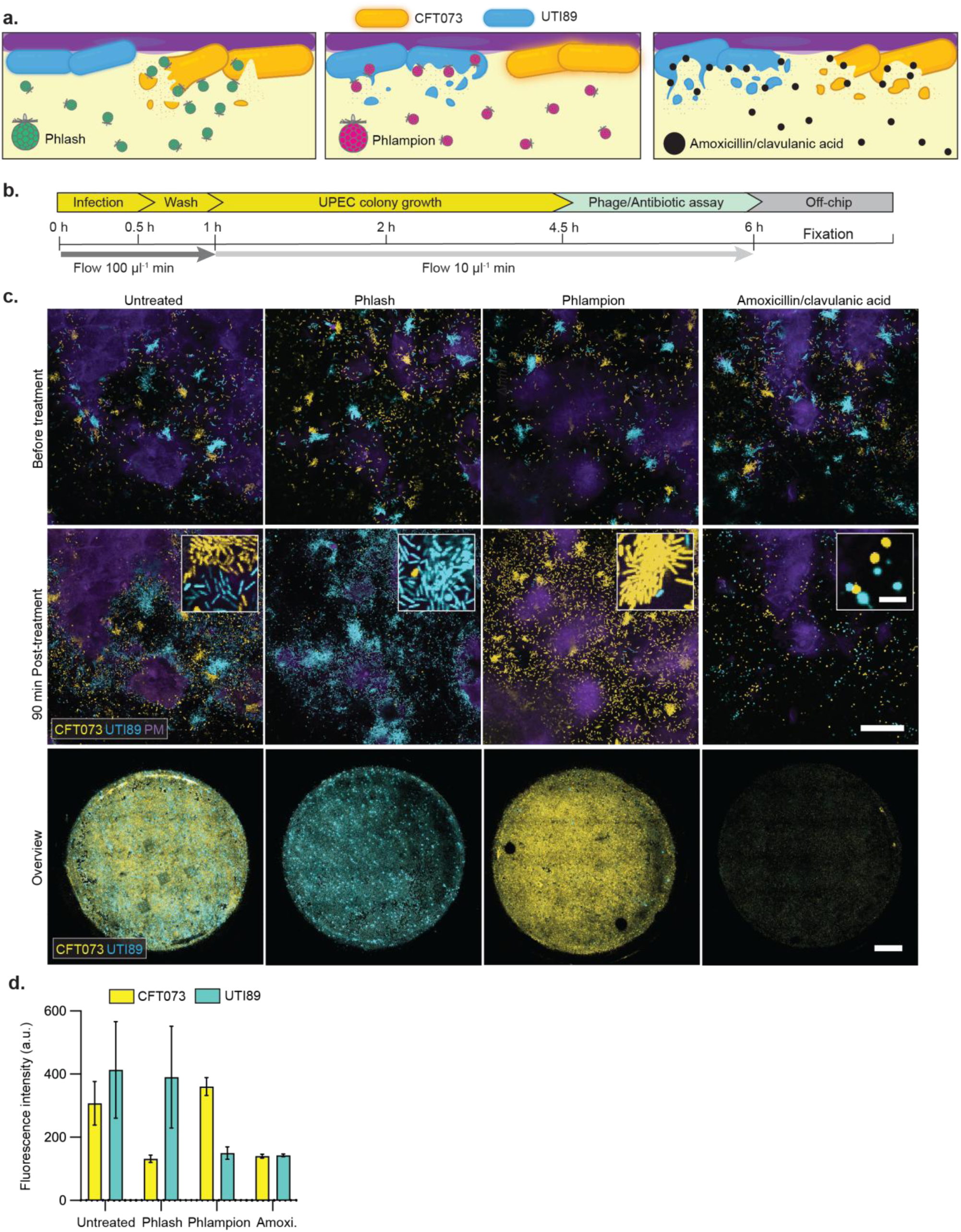
Targeted phage therapy selectively clears pathogenic strains from infected urothelium. **a**, Schematic illustration of treatments applied to the urothelial surface (plasma membrane, PM, purple) co-infected with UPEC strains CFT073 expressing mScarlet3 (yellow) and UTI89 expressing mGreenLantern (cyan). Infected tissues were treated with a CFT073-specific phage (Phlash, left), a UTI89-specific phage (Phlampion, middle), or the broad-spectrum antibiotic amoxicillin/clavulanic acid (right). **b**, Experimental workflow and corresponding timeline. **c**, The urothelium was co-infected with a 1:1 mixture of CFT073 expressing mScarlet3 (orange) and UTI89 expressing mGreenLantern (cyan) under continuous urine flow for 4.5 h, followed by 1.5h of treatment with Phlash, Phlampion, amoxicillin/clavulanic acid (amoxi), or no treatment (untreated). Top and middle rows: representative live-cell fluorescence images acquired before and after treatment (40× objective). Insets show close-ups after the respective treatments. Scale bars, 50 µm (main images) and 5 µm (insets). Bottom row: endpoint imaging of infected urothelium showing surface-associated CFT073 (yellow) and UTI89 (cyan). The images represent maximum intensity projections from stitched 10× fields. Data represent *n* = 3 biologically independent experiments with duplicates per condition. Scale bar, 1 mm. **d**, Quantification of CFT073 (yellow) and UTI89 (cyan) fluorescence intensities on the urothelial surface, corresponding to the overview images in (c). Data represent mean ± s.d. from *n* = 3 biologically independent experiments.

In untreated controls, both strains spread extensively across the tissue surface (Fig. 6c). Phage treatment resulted in selective clearance: phage Phlash eliminated CFT073, while phage Phlampion eradicated UTI89, leaving the non-targeted strain unaffected. In contrast, amoxicillin/clavulanic acid treatment targeted both strains, leaving only bacterial debris on the tissue surface (Fig. 6c). Endpoint imaging and fluorescence quantification confirmed the strain specificity of phage therapy and the broad efficacy of antibiotic treatment (Fig. 6d). Together, these findings demonstrate that our platform enables evaluation of both conventional (antibiotic) and unconventional (phage) therapies under physiologically relevant conditions, and highlight the potential of phage treatment as a selective, strain-specific alternative to broad-spectrum antibiotics.

## Discussion

In this study, we established a perfusable, three-dimensional (3D) human urothelium model integrated into a microfluidic platform that enables real-time, high-resolution live-cell imaging of urinary tract infections (UTIs) under physiologically relevant flow conditions. The system addresses two long-standing limitations of in vitro UTI models: (i) limited physiological fidelity, particularly with respect to native urothelial architecture and urine flow, and (ii) the inability to dynamically monitor infection and treatment responses over time and at high spatiotemporal resolution.

We first developed a robust method to generate a fully differentiated, urine-tolerant human urothelium on the underside of transwell inserts, allowing unobstructed optical access for high-resolution imaging. Building on previous upright transwell systems^12^, we introduced key modifications that supported inverted culturing and scalability to a 24-well format. The resulting tissue recapitulated the stratified architecture of the human bladder epithelium, including basal, intermediate, and umbrella cell layers that express physiologically relevant markers and maintain barrier function under urine exposure. To the best of our knowledge, this work represents the first demonstration of an inverted, urine-tolerant human urothelial microtissue that permits continuous live imaging under near-physiological conditions.

The integration of the inverted urothelium into a custom-designed microfluidic perfusion platform added a critical physiological dimension to the model. Urinary flow profoundly influences nutrient distribution, bacterial adhesion, and infection dynamics,^6^ but is absent from most current in vitro systems. Although the shear forces generated during micturition remain poorly defined in humans, flow is known to modulate the growth and organization of uropathogenic colonies.^20^ By reproducing this hydrodynamic environment, our platform provides a controllable and reproducible means to dissect how mechanical forces shape host-pathogen interactions. Previous efforts to introduce flow in other in vitro bladder models, such as the use of parallel plate flow chambers^21^ or in PDMS-based organ-on-chip devices^22^ typically required on-chip cell seeding, differentiation, and maintenance. Such workflows are technically demanding and offer limited compatibility with downstream tissue analysis. The modular design of our model decouples tissue maturation from subsequent infection and treatment assays, alleviating the technical and analytical constraints of on-chip differentiation. In parallel, the minimized optical working distance enhances both spatial and temporal resolution, enabling continuous visualization of infection progression and therapeutic responses at single-cell resolution.

Inverted configuration and dynamic perfusion proved critical to recapitulating the early stages of UPEC infection. The inverted orientation required active bacterial motility for tissue colonization, while urinary flow continuously removed planktonic cells and supplied nutrients to surface-associated bacteria, closely mimicking the dynamic equilibrium in the urinary tract. These features allowed us to resolve early host-pathogen interactions with unprecedented detail, including bacterial attachment, microcolony formation, and detachment under mechanical and pharmacological perturbation.

We benchmarked the system with clinical and ex vivo data for the antibiotic fosfomycin, which remains a first-line oral therapy for uncomplicated UTIs. Clinically, exposure to fosfomycin has been linked to the formation of bacterial L-forms, lacking a functional cell wall, in patient urine, with reversion to rod-shaped bacteria after drug removal as demonstrated ex vivo.^15^ Using controlled urine flow and exposure to clinically relevant antibiotic concentrations, our model faithfully recapitulated these dynamics: within hours of treatment, rod-shaped UPEC converted to L-forms that were stabilized during exposure to high osmolarity, and a small number of cells resumed growth after drug withdrawal. Importantly, high-resolution imaging in commercial CellASIC device under axenic conditions revealed that regrowth primarily originated from rod-shaped cells that were transiently non-dividing and metabolically quiescent, whereas L-form reversion to rod-shaped cells occurred only rarely. This finding offers a refined mechanistic explanation for UTI relapse after apparently successful antibiotic therapy and emphasizes the need to target non-dividing populations alongside L-form-mediated survival. The ability of our platform to reproduce clinically observed phenotypes provides strong validation of its predictive translational relevance.

We also demonstrated the translational utility of the developed platform by evaluating established and experimental therapeutic strategies. Real-time imaging revealed that D-mannose inhibited FimH-dependent bacterial adhesion and promoted detachment of existing microcolonies, consistent with its proposed mechanism as a competitive inhibitor of mannose-sensitive fimbrial binding. However, complete clearance was not achieved, probably due to persistent single bacteria interacting with the glycosaminoglycan-rich surface through FimH-independent mechanisms. This finding underscores the model’s sensitivity to subtle host and bacterial factors that shape infection persistence.

Finally, the system supported strain-specific bacteriophage therapy studies, in which targeted phages selectively eradicated UPEC colonies without affecting non-target strains, whereas broad-spectrum antibiotics, such as amoxicillin/ clavulanic acid, reduced all bacterial populations. Together, these data highlight the platform versatility for investigating both traditional and emerging antimicrobial strategies, including precision biologics.

Overall, this work establishes the first human-relevant microfluidic urothelial model that combines physiologically accurate tissue architecture, urinary flow, and live imaging to dissect UTI pathogenesis and therapeutic responses. By bridging the gap between static cultures and animal models, it provides a scalable, experimentally accessible platform for predictive preclinical testing of antibiotics, anti-adhesion agents, and phage therapies. Beyond UTIs, the modular design may be adapted to study other mucosal infections or drug delivery processes that involve dynamic epithelial barriers. Future work could incorporate immune and stromal components or patient-derived cells to extend the platform’s use in personalized infection modeling and the rational design of next-generation anti-infective strategies.

## Acknowledgements

Authors are grateful for the support of the Imaging Core Facility (IMCF) and the BioEM lab at Biozentrum, University of Basel, as well as of the Single Cell Facility (SCF) of D-BSSE, ETH Zürich. We thank Prof. Jennifer Rohn (University College London) for technical advice, Dr. Julie Sollier for helpful comments and Dr. Nichole Wespe for support in data management. This work was financially supported by the Swiss National Science Foundation (SNSF) within the framework of the National Competence Center in Research (NCCR) “AntiResist: New approaches to combat antibiotic-resistant bacteria”, under contract number 51NF40_180541.

## Author contributions

A.K. and I.S. conceived the study, designed the experiments, conducted experiments, analyzed and interpreted the data, and wrote the original manuscript draft. B.S. contributed to conceptualization and experiments, and analyzed and interpreted data. J.B. contributed to study conceptualization, data interpretation, and manuscript drafting. L.G. and M.U.G. optimized experimental protocols. L.G. and A.M. performed experiments. B.S., C.F., S.K. and S.T.-S. provided materials. C.D. and A.H. edited the manuscript, acquired funding, and supervised the project.

## Extended Data

**Extended Data Figure 1.**
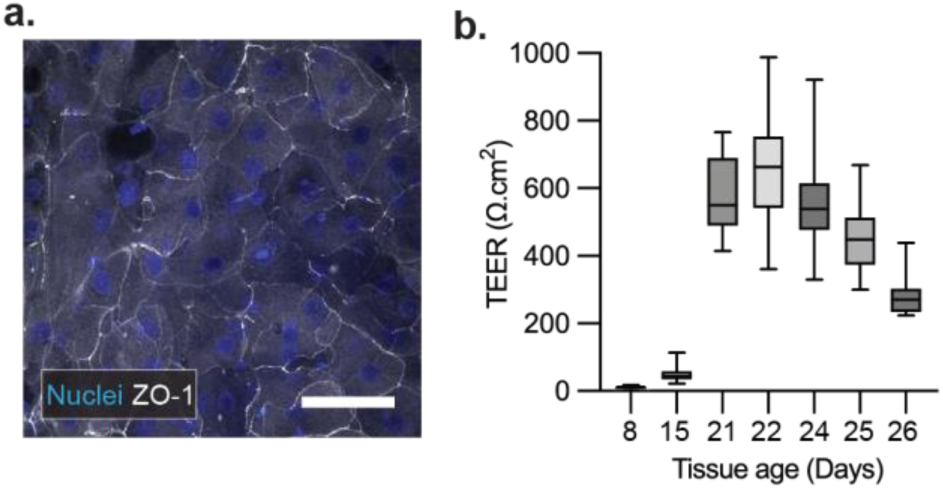
Tissue integrity and barrier function. **a**, Immunofluorescence analysis of the barrier-integrity marker ZO-1 (tight junction protein, white). The nuclei were stained with DAPI (blue). Scale bar: 20 µm. **b**, Transepithelial electrical resistance (TEER) measurements of tissues at the indicated time points during culturing, reflecting barrier maturation. Data are presented as box plots representing the mean ± s.d. of all inserts measured on each respective day. Owing to the longitudinal study design, the number of inserts varied across time points: Day 8 (n = 49), Day 15 (n = 46), Day 21 (n = 13), Day 22 (n = 88), Day24 (n = 45), Day 25 (n = 57), Day 26 (n = 10).

## Materials and Methods

### Strains, plasmids, and growth media

Targeted mutations in *Escherichia coli* CFT073 were generated using the λ Red recombination system. Expression was induced with 0.2% IPTG for 20 min. Linear DNA fragments encoding deletions in *fimH* or *fliC*, flanked by ∼40-bp homology arms and containing a kanamycin resistance cassette, were generated by PCR and purified. Electrocompetent CFT073(pWRG99) cells were electroporated with 2–8 µg of purified DNA (1.8 kV, 0.2-cm cuvette), recovered in SOC at 37 °C for 1 h, and plated on kanamycin-LB agar. Recombinants were screened by colony PCR and validated by Sanger sequencing. To cure pWRG99, strains were passaged at 42 °C and plated on non-selective media; plasmid loss was confirmed by antibiotic sensitivity and PCR.

Fluorescent *E. coli* strains were generated using plasmids encoding codon-optimized *mScarlet3* or *mGreenLantern* in a SC101-based vector under the control of a constitutive promoter. The strains were transformed by electroporation and selected on an LB agar plate supplemented with kanamycin (50 μg ml⁻¹).

### Bacterial cultivation

All strains were stored in glycerol at -80°C until further use. The CFT073, UTI89 and mutant uropathogenic *E. coli* strains were inoculated from an LB agar plate into LB medium and incubated overnight at 37°C. The culture was then diluted 1:50 in fresh LB medium and incubated for an additional 48 hours at 37°C. Antibiotic selection was applied throughout these steps to maintain plasmid expression. On the day of infection, the stationary-phase culture was diluted in pooled human urine to an optical density OD600 of 0.2, without antibiotic selection, and incubated with agitation at 150 rpm for 2 hours at 37°C until the culture reached the exponential phase. The bacteria cultures were then further diluted in urine to the desired OD600 values of 0.005–0.02. For simultaneous infection with two strains, the cultures were mixed in a 1:1 ratio, resulting in a final OD600 of 0.01. Colony-forming units (CFUs) for infection were determined by plating serial dilutions of the infection mix onto LA agar plates with antibiotic selection. The plates were incubated overnight at 37°C.

### Comparison of transwell infection setups

Infection assays were performed using either an i) upright-mimicking, ii) inverted static, and iii) inverted perfused setup.

i) For the upright-mimicking setup, the medium and urine were removed from both the upper and lower chambers of the transwell. The transwell was then flipped and placed onto the lid of a well plate. A 0.04 ml drop of the infection mix was added to the tissue. The infected transwell was then covered with the plate to avoid evaporation.

ii) For infections in the inverted static setup, the urine in the lower chamber of the transwell was replaced with 0.4 ml of the bacterial-urine infection mix.

In both orientations (i, ii), the infected tissue was incubated for 30 minutes at 37°C. After this incubation, the infection mix was directly removed, and 0.15 ml of 3D differentiation medium was added to the upper chamber, and 0.4 ml of urine was added to the lower chamber. The urine in the lower chamber was again removed, and the transwell was transferred to a new well, where 0.4 ml of fresh urine was added. The plate was then incubated for 2.5 h at 37°C. All incubation steps were carried out in a humidified incubator with 5% CO2.

iii) For infections in the inverted perfused setup, the infection mix was perfused through the microfluidic channels at a rate of 100 μl min^-1^ for 30 minutes. This procedure was followed by a 30-minute wash with fresh urine at the same flow rate to remove non-adherent bacteria. The flow rate was then reduced to 10 μl min^-1^ and maintained for 2 hours.

### Urine collection

Urine was collected anonymously from healthy volunteers of both sexes (> 250 donations). Donors with diabetes mellitus, antibiotic treatment within the previous 14 days, known pregnancy, menstrual bleeding on the day of sample collection, or current urinary tract infection were instructed not to donate urine, as stated in the written notice at the sampling sites. Individual urine samples were analyzed using a Combur9 dipstick test (Roche, Basel, Switzerland). Samples with elevated glucose levels or a volume of less than 50 ml were discarded. The female and male urine were pooled separately and sterile filtered. For experiments, the female and male urine pools were mixed in a 1:1 ratio, here designed as pooled human urine.

### Cell culture

HBLAK human bladder progenitor cells were purchased from CELLnTEC (Switzerland) as frozen vials (> 5×10^5^ cells ml^-1^). For the preparation of the working stocks, the thawed cells were mixed with prewarmed 2D growth medium (CnT-Prime, CELLnTEC, Switzerland) in a ratio of 1:10 in a 50 ml tube. Afterwards, the cells were harvested by centrifugation (300 g / 5 min / RT) and resuspended in a vented flask (75 cm^2^) with fresh 2D growth medium. The cells were cultivated at 37°C with 5% CO_2_ to 80–90 % confluence before being passaged. The cells of passage 4 were then frozen in a defined freezing medium (CnT-CRYO-50, CELLnTEC, Switzerland) at a concentration of 5×10^5^ cells ml^-1^ and stored in liquid nitrogen.

Cells routinely grown in 2D growth medium were sub-cultivated twice a week at a seeding density of 5×10^5^ cells per T75 cm^2^ flask using Accutase (CnT-Accutase-100, CELLnTEC, Switzerland) for cell detachment after washing with Ca^2+^ and Mg^2+^-free phosphate-buffered saline (PBS, Bioconcept, Allschwil, Switzerland).

### Tissue generation

HBLAK cells of passage 10 or 12 were used for seeding on Corning® 6.5 mm transwells with 0.4-µm pore size polycarbonate membrane (Corning 3413, Costar, NY, USA) using a suspension of 2.5×10^6^ cells ml^-1^. All incubation steps were carried out at 37°C with 5% CO_2_ in a humidified incubator. Prior to seeding, the 24-well plate was flipped, positioning the transwells on the lid with the bottom side of the membrane facing upward. The cell suspension was then distributed onto the membrane, ensuring complete coverage. The transwells were covered with the well plate to prevent evaporation, with a spacer inserted between the lid and the plate to avoid contact between the cell suspension and the surface of the plate.

Following a 4-hour incubation at 37°C with 5% CO2 in a humidified incubator, 50 µl of 2D growth medium (CnT-Prime, CELLnTEC, Switzerland) were added to the seeded cells. After an additional 16 hours of incubation at 37°C, the well plate was flipped to its original orientation and 2D growth medium was added to both sides of the transwell insert (0.1 ml upper chamber, 0.6 ml lower chamber) on day 1. To validate tissue development, Invitrogen CellMask™ Deep Red Plasma Membrane Stain (C10046, Thermo Fisher, Waltham, MA, USA) was added to the medium (dilution 1:5000). On the fourth day, when the cells had reached confluence, the medium was changed (0.1 ml and 0.6 ml) to 3D differentiation medium (CnT-PR-3D, CELLnTEC, Switzerland), which had been supplemented with growth factor mix (00192152, KGM Gold SingleQuots, Lonza, Basel, Switzerland). During the cell culture and the generation of tissue, no antibiotics were added. Subsequent medium changes were executed on days 6, 8, 11 and 13. On day 15, tissues were placed at an air-liquid interface (ALI), with the medium aspirated from both chambers of the transwell and 0.15 ml of 3D differentiation medium being added to the upper chamber. Cells were maintained under ALI conditions until day 22 with additional medium changes occurring on days 18 and 20. On day 22, the tissues were exposed to a urine-liquid-interface (ULI) by adding 0.15 ml of 3D-Prime medium to the upper chamber and 0.4 ml of pooled human urine to the lower chamber. Subsequent medium and urine exchanges were carried out at 2-day intervals until the conclusion of the experiments. The experiments were carried out between days 24 and 26.

### TEER measurements

The transepithelial electrical resistance (TEER) of the tissues was measured using an EVOM volt–ohm resistance meter (World Precision Instruments, Berlin, Germany), equipped with STX4 chopstick electrodes (World Precision Instruments). For TEER measurements, the inserts were submerged in 37 °C warm HBSS (300 µl upper chamber and 1 ml lower chamber). From each endpoint measurement point, the TEER value of an empty transwell was subtracted and normalized to the surface area of the membrane (0.33 cm^2^).

### Immunocytochemistry

Prior to immunofluorescence staining, tissues were fixed overnight at 4°C in 4% paraformaldehyde (PFA; 15710, Electron Microscopy Sciences, Hatfield, PA, USA) prepared in HBSS. After fixation, membranes were excised using a scalpel, transferred to a 24-well plate, and permeabilized in HBSS containing 0.2% Triton X-100 (Sigma-Aldrich, St. Louis, MO, USA) for 20 minutes at room temperature (RT). After a single wash with HBSS, samples were blocked for 1 hour at RT in HBSS, supplemented with 1% normal goat serum (50062Z, Thermo Fisher Scientific, Waltham, MA, USA) and 1% bovine serum albumin (BSA; A9647, Sigma-Aldrich, St. Louis, MO, USA). The tissue-containing membranes were then incubated overnight at 4° C with primary antibodies diluted in HBSS containing 1% BSA. The following primary antibodies (all at 1:100 dilution) were used: mouse anti-uroplakin-IIIa (UPKIIIa) monoclonal antibody (sc-166808, Santa Cruz Biotechnology, Dallas, TX, USA), rat anti-heparan sulfate proteoglycan 2 monoclonal antibody (ab2501, Abcam, Cambridge, UK), and mouse anti-cytokeratin 5 monoclonal antibody (MA5-17057, Thermo Fisher Scientific, Waltham, MA, USA). Following incubation with primary antibodies, the membranes were washed and incubated for 1.5 hours at room temperature (RT) with secondary antibodies diluted 1:200 in HBSS containing 1% BSA. The following Alexa Fluor 488-conjugated secondary antibodies were used: goat anti-mouse (A-11001) and goat anti-rat (A-11006), both from Thermo Fisher Scientific (Waltham, MA, USA). CellMask™ Deep Red Plasma Membrane Stain (1:5000 dilution; C10045, Thermo Fisher Scientific, Waltham, MA, USA) and DAPI (1 mg ml^-1^; D1306, Thermo Fisher Scientific, Waltham, MA, USA) were added concurrently to the secondary antibody solution for counterstaining. After staining, the membranes were washed, mounted using ProLong™ Glass Antifade Mountant (P36980, Thermo Fisher Scientific, Waltham, MA, USA), and imaged using a Nikon CSU W1 spinning disk confocal microscope. Images were acquired at 40×, 60×, or 100× magnification.

For the analysis of tight junctions, no blocking step was performed. The tissues were incubated at RT for 3.5 hours in HBSS with the following reagents: Alexa Fluor 555-conjugated anti-zonula occludens-1 (ZO-1) monoclonal antibody (1:150; MA3-39100-A555), DAPI (1 mg ml^-1^; D1306), and CellMask™ Deep Red Plasma Membrane Stain (1:5000 dilution; C10045), all reagents were purchased from Thermo Fisher Scientific (Waltham, MA, USA).

### Microfluidic platform fabrication

Poly(methyl methacrylate) (PMMA)-based microfluidic devices (25 mm × 75 mm) were fabricated using a computer numerical control (CNC) micro-milling system (DATRON Neo, DATRON Dynamics, Milford, NH, USA), as detailed previously. Each chip contained four identical micro-milled channels, each with a height of 300 µm and featuring a central open well designed for inserting cell-coated transwells. The channels were sealed at the bottom using glass coverslips (ref. 10812, ibidi GmbH, Martinsried, Germany) secured with a biocompatible epoxy adhesive (MED-302-3 m, Epoxy Technology Europe AG, Cham, Switzerland).

Each insertion well was equipped with a groove slot (height: 2.4 mm, width: 2.2 mm) fitted with an elastic O-ring (8.6 × 2.4 mm, item no. 860400.0258, Brütsch/Rüegger Werkzeuge AG, Urdorf, Switzerland).

To minimize contamination risk and reduce media evaporation, the basolateral side of each transwell was closed with a custom-designed lid manufactured from GreenTec Pro material. Each channel featured inlet and outlet openings (diameter: 0.8 mm each) to facilitate liquid perfusion.

Following assembly, the microfluidic chips were sterilized by washing with 50% ethanol, followed by UV exposure on both sides for 30 minutes. After sterilization, each channel was coated with 200 µl of BIOFLOAT FLEX coating solution (F202005, Facellitate, Mannheim, Germany) and incubated for 10 minutes at room temperature to minimize non-specific bacterial adhesion to the channel walls and glass coverslip during subsequent infection assays.

### Transition from the standard culture to the perfused culture

Before insertion in the chip, the contents of both basal (3D differentiation medium) and apical (urine) compartments of the transwell inserts were fully aspirated. The transwell was manually positioned above the insertion well and carefully inserted through the elastic O-ring. Insertion expanded the O-ring laterally, which then tightly sealed around the transwell insert to prevent leakage. Once the transwell inserts were fully inserted, the channels were manually filled with 50 µl of warm urine using a P200 pipette. The basal compartment was filled with 150 µl of 3D differentiation medium. The microfluidic chip with the inserted transwells was then transported to the microscope and connected to the fluidic setup to perform the real-time imaging assay.

### Fluidic setup assembly for infection and therapeutic assay

The microfluidic platform was connected to the fluidic circuit through the corresponding inlet and outlet connectors. The chip was placed inside a microscope stage incubator at 37°C in an atmosphere of 5% CO2 and 95% relative humidity.

A peristaltic pump (ref. 78001–18, Ismatec, Switzerland) was used to perfuse urine into the chip. Upstream, peristaltic tubings (070534-08L-ND, ID 0.76 mm, wall 0.85 mm, Idex Health & Science GmbH, Wertheim, Germany) were connected to 50-cm-long polytetrafluoroethylene (PTFE) tubing (ID 0.3 mm, OD 0.6 mm, Bola GmbH, Grünsfeld, Germany) through metal connector pieces, obtained from standard Luer-lock syringe-tubing connectors (IP721-90, GONANO Dosiertechnik GmbH, Breitstetten, Austria). These pieces were connected to a long needle that went into a falcon tube containing urine. Downstream, peristaltic tubing was connected to 100-cm-long PTFE tubing that was connected to the inlet of the chip through a 2-cm-long tygon-tube (ID 0.76 mm) connector. The chip outlet was connected to a collection reservoir through 50 cm polytetrafluoroethylene (PTFE) tubing (ID 0.5 mm, OD 1.0 mm, Bola GmbH, Grünsfeld, Germany) through metal connecting pieces, obtained from standard Luer lock syringe tubing connectors (TE 27 GA 90◦ bent, APM Technica AG, Heerbrugg, Switzerland). The inflow rate of the peristaltic pumps was set to either 10 µl min^−1^ or 100 µl min^−1^ according to the different steps of the assay.

### Fluorescent nanoparticle perfusion test

Carboxyl fluorescent particles (0.46 µm) (GFP-0552-2, Spherotech, Lake Forest, IL, USA) were diluted in warm urine (1:2000) and perfused in the microfluidic channel at 100 µl min^−1^. Live movies of particle flow underneath the tissue model were acquired using the same microscope setup as in the Infection and Assay sections.

### CellASIC microfluidic assays

Bacteria were loaded into B04A CellASIC® ONIX microfluidic plates (Merck Millipore) following the manufacturer’s instructions. Cultures were prepared as described for the tissue infection experiments. Cells were trapped within the microfluidic growth chambers under controlled pressure (7 kPa) using the ONIX microfluidic system, which maintained continuous media perfusion throughout the experiment. Bacteria were first incubated for 3 h in individual ex vivo filter-sterilized urine samples, followed by a 5 h treatment with 400 mg L⁻¹ fosfomycin in the same samples. After treatment, the channels were perfused with urine lacking fosfomycin. Time-lapse imaging was performed at 37°C on a Nikon Eclipse Ti2 inverted fluorescence microscope equipped with a 100× oil immersion objective (CFI Plan Apo λDM 100× Oil, Nikon) to monitor single-cell growth and morphology over time. Exposure settings were 100 ms for phase contrast, and 50 ms (15% light intensity) for both mGreenLantern and mScarlet3 fluorescence channels.

### Time-kill curves

Bacterial time-kill experiments were conducted in sterile 96-well microtiter plates. *E. coli* CFT073 cultures, prepared as described for the tissue infection experiments, were diluted in individual ex vivo filter-sterilized urine samples to achieve an initial inoculum of approximately 10⁶ CFU/mL per well. Fosfomycin was added at a final concentration of 400 mg L⁻¹, and plates were incubated statically at 37°C. Starting from the time of antibiotic addition, samples were collected hourly, serially diluted tenfold in phosphate-buffered saline, and plated on LB agar. Colony-forming units (CFU) were enumerated after incubation at 37°C for 16 h.

### Fosfomycin treatment

An uropathogenic Escherichia coli (UPEC) strain CFT073 (wild type, WT) constitutively expressing a fluorescent reporter was used to monitor antibiotic response dynamics in real time. Overnight cultures were grown in LB medium at 37 °C with shaking, then subcultured 1:100 into fresh medium and incubated to mid-log phase (OD600 ∼0.4-0.6). Bacteria were harvested by centrifugation, washed once in sterile phosphate-buffered saline (PBS), and resuspended in human urine to a final concentration of approximately 10⁷ CFU ml⁻¹. The bacterial suspension was perfused through the microfluidic channels at a rate of 100 μl min⁻¹ for 30 min to initiate infection, followed by a 30 min wash with pooled sterile urine at the same flow rate to remove non-adherent bacteria.

Subsequently, the flow rate was reduced to 10 μl min⁻¹ and maintained for 2 h to allow microcolony formation on the urothelial surface. Fosfomycin treatment was then initiated by perfusing urine samples of different osmolarities, two high (M1, F2) and two low (F3, F6), supplemented with fosfomycin at a final concentration of 400 mg l⁻¹. The treatment lasted 5 h under continuous perfusion at 10 μl min⁻¹, during which time-lapse imaging was performed every 30 min using a 10× objective to monitor the change on the tissue-pathogen interface. Following treatment, the perfusate was replaced with antibiotic-free urine of the corresponding samples and flushed through the channels at 100 μl min⁻¹ for 30 min to remove residual fosfomycin. The flow was then stopped, and imaging was continued using a 40× objective to capture time-lapse sequences of UPEC regrowth from surviving L-form cells.

The effect of fosfomycin treatment was quantified from 3 independent experiments (n = 3). In each experiment, two technical replicates (transwell inserts) were analyzed per condition, with three regions of interest (ROIs) imaged per insert at 10× magnification and time point. UPEC surface coverage was quantified in Fiji (ImageJ) by Triangle-method thresholding, and area fraction values were extracted and normalized to baseline (T0 = 100%). For each experiment, values represent the mean of three ROIs per insert, averaged across duplicate inserts, and subsequently combined to obtain the grand mean across experiments.

### D-mannose assay

An uropathogenic *Escherichia coli* (UPEC) strain CFT073 (wild type, WT) and its isogenic Δ*fimH* mutant were used for mannose inhibition assays. Both strains were engineered to constitutively express fluorescent markers to enable live imaging. Overnight cultures were grown in LB medium at 37 °C with shaking, then subcultured 1:100 into fresh medium and incubated to mid-log phase (OD600 ∼0.4-0.6). Bacteria were harvested by centrifugation, washed once in sterile phosphate-buffered saline (PBS), and resuspended in diluted human urine to a final concentration of approximately 10⁷ CFU ml^-1^. The bacterial suspension was perfused through the microfluidic channels at a rate of 100 μl min^-1^ for 30 minutes to initiate infection. This procedure was followed by a 30-minute wash with sterile urine at the same flow rate to remove non-adherent bacteria. The flow rate was then reduced to 10 μl min^-1^ and maintained for 1 hour. Beginning at 2 hours post-infection (p.i.), tissues were perfused with either sterile urine (control) or urine, supplemented with D-mannose (Sigma-Aldrich, St. Louis, MO, USA) at final concentrations of 1% or 7% (w/v). The assay continued until 5 hours p.i., during which high-resolution live imaging was performed to monitor the dynamics of mannose-mediated clearance of the infection. At the endpoint, tissues were retrieved from the device and fixed for subsequent imaging and analysis. To evaluate mannose-mediated colony clearance, bacterial colonization was quantified by measuring residual fluorescence intensity on the tissue surface following treatment with three mannose concentrations (0%, 1%, and 7%). Whole-tissue images were acquired at 10× magnification, containing both the wild type and Δ*fimH* mutant strains. Maximum-intensity projections were generated in Fiji (ImageJ), then analyzed by calculating the fluorescence intensity (“Mean Intensity” function) within a standardized circular region of interest (ROI; 6.0-mm diameter). Data are presented as mean ± s.d. from three independent experiments (n = 3).

### Bacteriophage isolation

Bacteriophages were isolated from freshwater samples of the Rhine river in Basel, Switzerland. The isolation procedure followed the method described by Maffei et al.^23^, with the addition of an enrichment step using CFT073 or UTI89 to increase phage yield prior to plaque isolation. Briefly, freshwater samples containing phages were enriched by adding a 1:5 dilution of 5× LB without salt, followed by the addition of CFT073 or UTI89 from an overnight culture at a 1:10’000 dilution. The mixture was incubated overnight, after which 0.1 ml was plated for a plaque assay using the double-agar overlay method.^24^ The assay involved 25 ml of LB agar as base and 4 ml of LB top agar containing 0.4% agar as the overlay in round Petri dishes. After adding 0.1 ml of the bacterial overnight culture to the top agar, the plate was incubated at 37°C for at least 12 hours. Once the plaques appeared, they were streaked at least three times on plates containing fresh agar and prey-bacteria to isolate single plaques.

### Bacteriophage storage

A high-titer stock of bacteriophages was prepared using the double-agar overlay method to achieve semi-confluent plaque coverage on the bacterial lawn. Following this step, 12 ml of SM buffer was added to the plate, and the suspension was gently agitated for at least 48 hours at 4°C. The suspension was then centrifuged at 8,000 *g* for 10 minutes in 15 ml tubes containing 0.35 ml of chloroform. The resulting supernatant, containing the phages, was transferred to glass vials and stored at 4°C.

### Phage assay

CFT073 and UTI89 UPEC strains, each expressing distinct fluorescent markers, were used for co-infection assays. Overnight cultures were grown in LB at 37 °C, subcultured to mid-log phase, washed, and resuspended in diluted human urine to a combined concentration of ∼10⁷ CFU ml^-1^ (1:1 ratio). The bacterial mixture was perfused into the microchannels at 100 μl min^-1^ for 30 minutes, followed by a 30-minute wash with sterile urine. The flow was then reduced to 10 μl min^-1^ and maintained for 3 hours to allow widespread colonization of the urothelium. At 4 hours post-infection (p.i.), tissues were perfused with either sterile urine (control), urine containing Phlash (targeting CFT073), urine containing Phlampion (targeting UTI89), or urine supplemented with 3.2 μg ml^-1^ amoxicillin/clavulanic acid. Treatments continued until 5.5 h p.i., during which high spatial-temporal-resolution imaging was performed to monitor treatment-specific dynamics. The tissues were then fixed and imaged for the endpoint analysis of the remaining bacterial burden. To evaluate treatment efficacy, bacterial clearance was quantified by measuring residual fluorescence on the tissue surface. Following treatment, whole-tissue images were acquired at 10× magnification. Maximum-intensity projections were generated in Fiji (ImageJ), and fluorescence intensity was quantified using the “Mean Intensity” function within a standardized circular region of interest (ROI; 6.0 mm diameter). Data are presented as mean ± s.d. from three independent experiments (n = 3). For each independent experiment, values were obtained by averaging available technical replicates (1–2 inserts) per condition. Fluorescence signal was measured for both CFT073 and UTI89 strains.

### Numerical simulations

The fluid flow in the platform was modeled in COMSOL (COMSOL Multiphysics 6.0, COMSOL Multiphysics GmbH, Zürich, Switzerland) using the Laminar Flow module.

### Image analysis

The images were analyzed using NIS-Elements General Analysis software, FIJI ImageJ (v1.54d), and Imaris Imaging Analysis software.

### Statistical analysis

Statistical analysis was performed using GraphPad Prism. All results are shown as mean values ± SD.

