## Supplementary information for "Perfusable 3D human urothelial model for real-time analysis of bacterial infection dynamics and therapeutic interventions"

for the manuscript

by **Kurmashev, A., Sorg, I. *et al.***

a

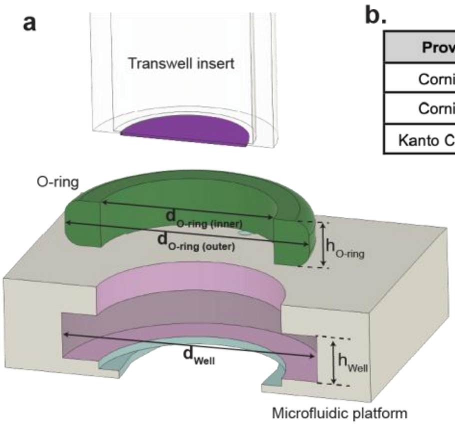


b. Adaptations of the microfluidic platform for three commerical Transwell inserts

| Provider | Product № | Porous membrane | $d_{well}$ | $h_{well}$ | $d_{O\text{-}ring(outer)}$ | $d_{O\text{-}ring(inner)}$ | $h_{O\text{-}ring}$ |
|---|---|---|---|---|---|---|---|
| Corning ® | 3413 | Polycarbonate | 14 | 2.4 | 13.4 | 8.6 | 2.4 |
| Corning ® | 3470 | Polyethylene terephthalate | 13.2 | 1.75 | 12.29 | 8.73 | 1.78 |
| Kanto Chemical | 08364-96 | Collagen Vitrigel™ | 13.1 | 2 | 13 | 9 | 2 |

c.

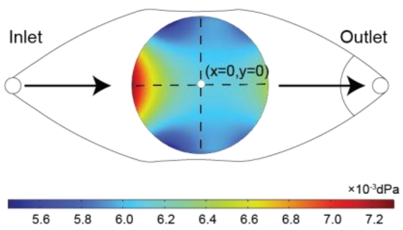


d.

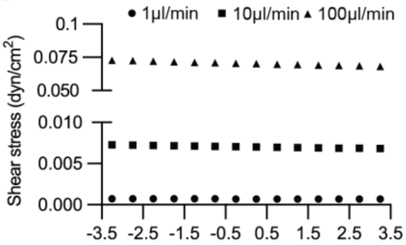


**Supplementary Figure S1. Design and Flow Simulation of the Microfluidic Platform**

**a,** Schematic illustrating the insertion of a transwell into the microfluidic unit, incorporating an O-ring to ensure tight sealing. **b,** Engineering modifications to the O-ring dimensions and the receiving well to enable compatibility with three standard commercial transwell formats. **c,** COMSOL model showing shear stress across the tissue surface during urine perfusion at 100 µl $min^{-1}$. **d,** Shear stress distribution along the flow direction (cross-section indicated in a) is shown for flow rates of 1, 10, and 100 µl $min^{-1}$.

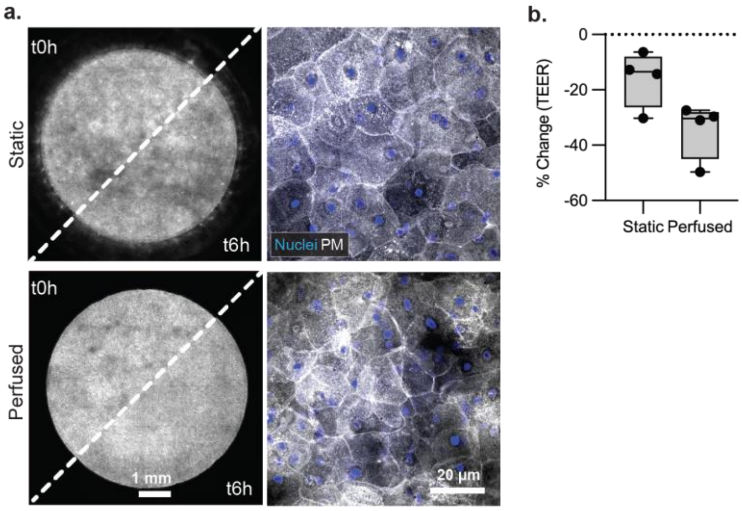


**Supplementary Figure S2. Structural and functional integrity of uroepithelium under static and flow conditions.**

**a,** Live 10× overview images of the uroepithelium at the start (t0h) and after 6 hours (t6h) of incubation under static conditions (top left) or 100 µl min$^{-1}$ urine perfusion (bottom left), with each image split diagonally to show the states before and after. Scale bar: 1 mm. Right panel: 40× endpoint images of fixed tissues (t6h) stained for nuclei (cyan) and plasma membrane (purple, PM), evidencing preserved umbrella cell architecture under both conditions. Scale bar: 50 µm. **c,** Relative change in TEER after 6 h static or perfused urine incubation, normalized to baseline (t0h). Barrier integrity declined under both conditions, with a larger drop under perfusion.

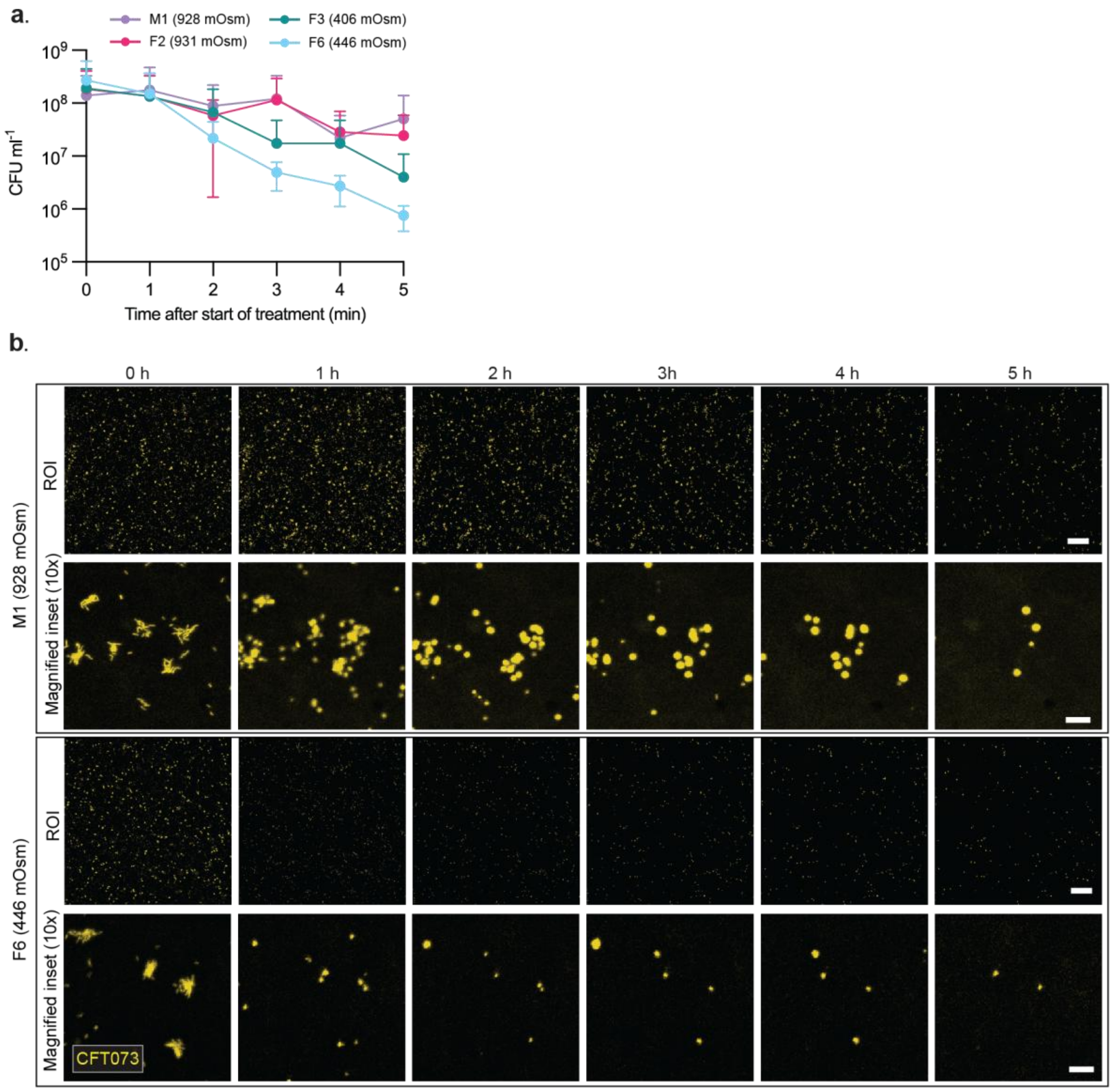


**Supplementary Figure S3. Osmolarity-associated fosfomycin response in urine and in the urothelial tissue model. a,** Time–kill curves of UPEC strain CFT073 expressing mScarlet3 (yellow) cultured in urine samples of high (M1, F2) and low (F3, F6) osmolarity and exposed to 400 mg $l^{-1}$ fosfomycin under conventional batch-culture conditions. **b,** Representative fluorescence images showing bacteria on the urothelial tissue surface during fosfomycin treatment under continuous urine flow. Images correspond to the high-osmolarity sample M1 and low-osmolarity sample F6 and were used to generate the quantitative analysis shown in Fig. 4b. The top row shows wide-field views, and the bottom row shows 10× magnified insets highlighting local bacterial aggregates and lysis dynamics over time. Scale bars, 200 µm (top) and 20 µm (bottom).

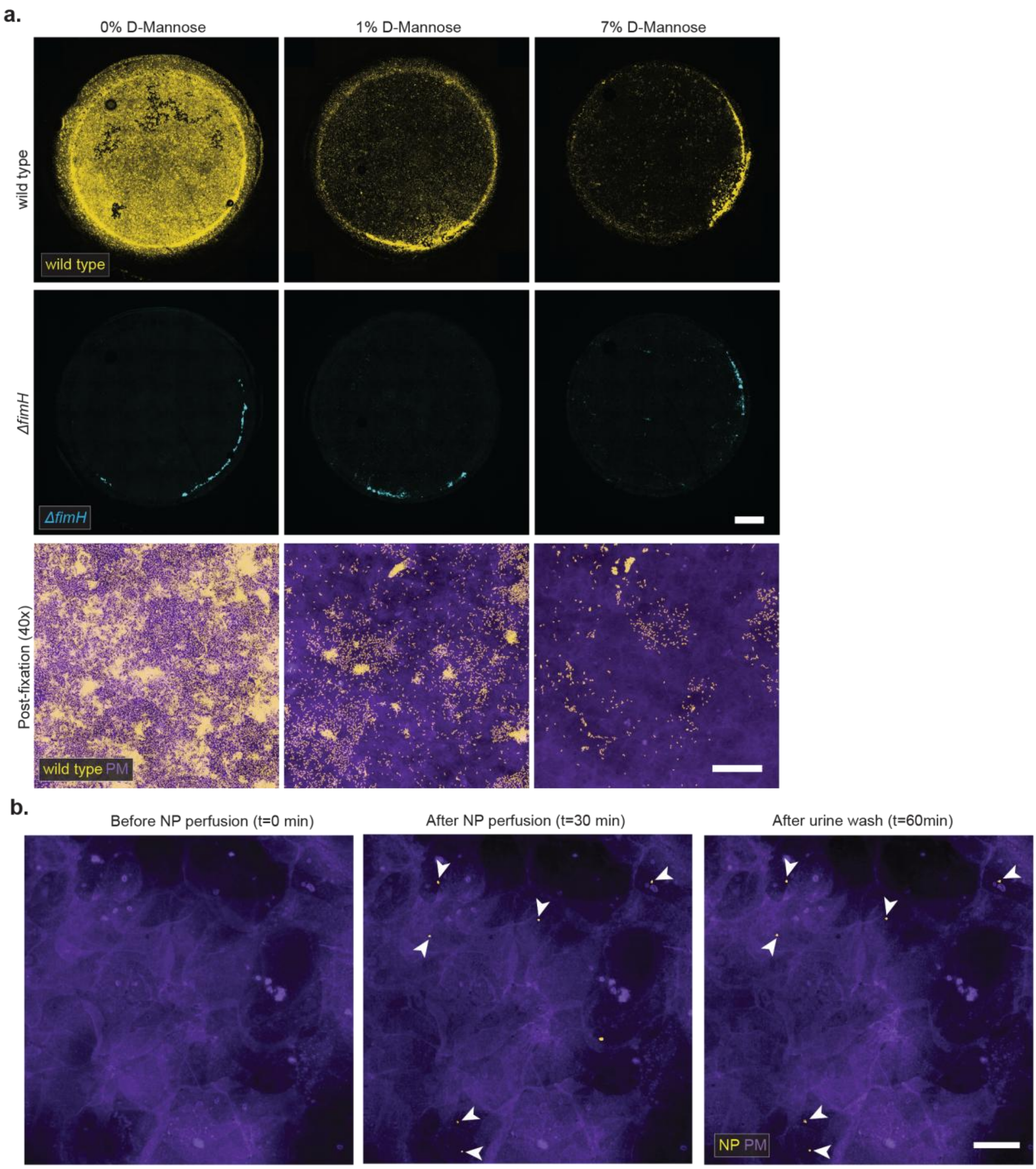


**Supplementary Figure S4. D-mannose–dependent inhibition of UPEC colonization.**

**a,** Fixed-tissue endpoint images showing urothelial colonization by UPEC strain CFT073 expressing mScarlet3 (wild type, yellow) and the isogenic mutant Δ*fimH* expressing mGreenLantern after treatment with 0%, 1%, or 7% D-mannose. 10× overview images showing reduced surface-associated wild type with increasing mannose concentration and uniformly low signal from Δ*fimH* UPEC across all conditions (upper and middle panels). Representative 40× images of wild type UPEC colonization (lower panel). Scale bars: 1 mm and 50 μm. **b,** Representative images showing the urothelium (left), after 30 min of nanoparticle perfusion in urine (middle), and after an additional 30 min flush with clear urine (right). Low-level adhesion of inert nanoparticles indicates a non-specific binding mechanism that may also account for residual *ΔfimH* UPEC attachment. Scale bar: 20 μm.

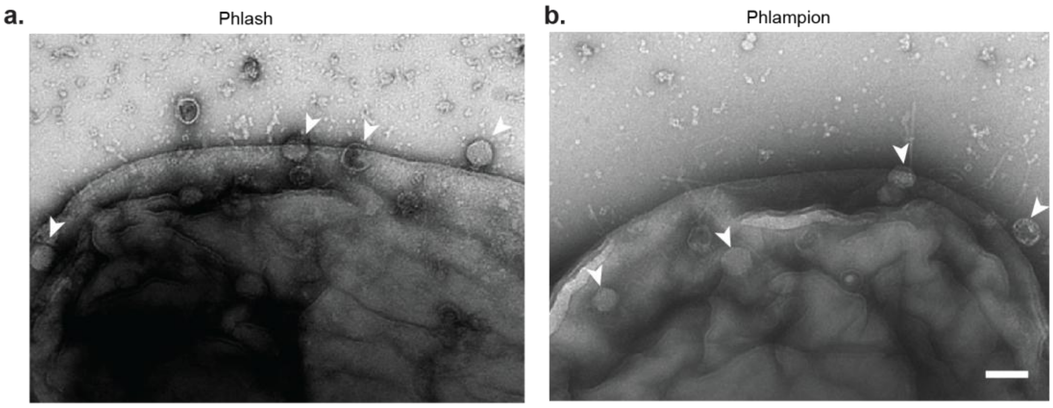


**Supplementary Figure S5. Electron microscopy images of strain-specific phages that interact with UPEC cells.**

**a**, Phlash in contact with the *E. coli* strain CFT073. **b**, Phlampion in contact with *E. coli* strain UTI89. Shown are transmission electron microscopy images negatively stained with 2% (w/v) uranyl acetate. Scale bars, 100 nm.

**Supplementary Table S1: Analysis of bacterial regrowth in urine in axenic CellASIC flow experiments**

| Urine ID | Osmolarity | Regrowth events from rods | Regrowth events from L-forms |
|---|---|---|---|
| M1 | 928 | 0 | 0 |
| M2 | 895 | 16 | 1 |
| F1 | 1025 | 1 | 0 |
| F2 | 931 | 12 | 0 |
| F3 | 406 | 2 | 0 |
| F4 | 219 | 2 | 0 |
| F6 | 446 | 11 | 0 |
| **Total** | | **44** | **1** |

Quantification of the number of bacteria with the ability to survive fosfomycin treatment and resume growth after antibiotic removal in urine samples. Data was acquired using the CellASIC microfluidics platform. Note that the number of regrowth events are not directly translatable to the protective effect of the individual urine samples due to differences in microfluidic cell loading.

**Supplementary Movie S1. Bacterial regrowth after fosfomycin washout by L-form reversion to rod-shape cells**

Live 100× imaging of CFT073 expressing mGreenLantern in CellASIC growth chambers for 17h. A continuous flow of urine (ID: M2) provided fresh nutrients to sustain growth. After 3 h, a dose of 400 mg $l^{-1}$ fosfomycin was administered for 5 h, as indicated. The video shows bacteria lysing due to the action of Fosfomycin and the emergence of cell-wall free L-form cells, which transiently revert to the rod-shape after antibiotic removal.

**Supplementary Movie S2. Bacterial regrowth after fosfomycin washout from rod-shaped bacteria that were transiently non-dividing and metabolically quiescent**

Live 100x imaging of CFT073 expressing mGreenLantern in CellASIC growth chambers for 17h. A continuous flow of urine (ID: F4) provided fresh nutrients to sustain growth. After 3 h, a dose of 400 mg $l^{-1}$ fosfomycin was administered for 5 h, as indicated. The video shows bacteria lysing due to the action of Fosfomycin and the survival of some cells with a dormant, non-dividing phenotype. These cells resumed growth after removal of the antibiotic, without transitioning through the L-form state.

**Supplementary Movie S3. Mannose-mediated detachment of UPEC colonies from the urothelium.**

Live 40× imaging of the urothelium at 5 h p.i. under perfusion of urine spiked with 1% D-mannose. The video shows (i) the detachment of a preformed WT UPEC microcolonies and (ii) the resulting sparse distribution of individual bacteria, mimicking the Δ*fimH* mutant phenotype (Fig. 4c).
